# Determine transposable genes when the orders of genes are different

**DOI:** 10.1101/2023.03.14.532623

**Authors:** Yue Wang

## Abstract

Certain nucleotide sequences in DNA can change their positions. Such nucleotide sequences might be shorter than a general gene. When we restrict to nucleotide sequences that form complete genes, we can still find genes that change their relative locations in a genome. Thus for different individuals of the same species, the orders of genes might be different. Such spatial difference of gene orders might be affected by temporal difference of gene (mutation) orders, and can be used to explain the order of mutation problem in myeloproliferative neoplasm. A practical problem is to determine such transposable genes in given gene sequences. Through an intuitive rule, we transform the biological problem of determining transposable genes into a rigorous mathematical problem of determining the longest common subsequence. Given several number sequences, determining the longest common subsequence is a classical problem in computer science. Depending on whether the gene sequence is linear or circular, and whether genes have multiple copies, we classify the problem of determining transposable genes into different scenarios and design corresponding algorithms. Specifically, we study the situation where the longest common subsequence is not unique.

## 1 Introduction

Various events, such as inversion, insertion, deletion, and duplication, can change the nucleotide sequence [28]. Such rearrangement events lead to the existence of transposons (also called transposable elements or jumping genes), which are DNA sequences that can change their relative positions within the genome. Transposons were first discovered in maize by Barbara McClintock [41]. Transposons have various types: long terminal repeats (LTR) retrotransposons, Dictyostelium intermediate repeat sequence (DIRS)-like elements, Penelope-like elements (PLE), long interspersed elements (LINE), short interspersed elements (SINE), terminal inverted repeats (TIR), Helitrons, etc. [40].

Transposons are common in various species. For the human genome, the proportion of transposons is approximately 44%, although most of transposons are inactive [42]. Transposons can participate in controlling gene expression [82], and they are related to several diseases, such as cancer [13], hemophilia [33], and porphyria [45]. Transposons can drive rapid phenotypic variations, which cause complicated cell behaviors [80, 48, 47, 11, 29]. Transposons can be used to detect cancer drivers [49] and potential therapies [2]. Transposons are also essential for the development of *Oxytricha trifallax* [50], antibiotic resistance of bacteria [3], and the proliferation of various cells [54, 78, 14]. With the presence of transposons, the regulation between genes might be affected, which is a challenge for inferring the structures of gene regulatory networks [74] and general transcriptome analysis [60, 81].

When transposons have been determined, we can use them to compare the genomes of different species, and such comparisons can be combined with other measurements between species, such as metrics on developmental trees [70]. Such comparisons can be also extended to different tissues to help with the prediction of tissue transplantation experiments [75]. Besides, for some species, cells at different positions have different gene expression patterns, which might be related to transposons [72].

Many transposons are as short as 10^2^ – 10^3^ base pairs, shorter than a general gene [53]. To determine such short transposons, one needs to analyze the original AGCT nucleotide sequences. There have been many algorithms developed to determine short transposons from nucleotide sequences, such as MELT (Mobile Element Locator Tool) [18], ERVcaller (Endogenous Retro-Virus caller) [10], and TEMP2 (Transposable Elements Movements Present 2) [79]. Different algorithms may only determine certain types of transposons. For more details, readers may refer to other papers [51, 20]. They use raw DNA sequencing data, which only contain imperfect information about the true DNA sequence, and the data quality depends on some factors that vary across different datasets [17]. Besides, they need a corresponding genome or reference transposon libraries.

There are gross DNA changes that associate with many genes, also called genomic rearrangements [21]. Such rearrangements include inversion, transposition, fusion, and fission [8]. To determine such gross genomic rearrangements, one first needs to convert nucleotide sequences into gene sequences by annotation. For two different gene sequences, the general idea of determining rearrangements is to calculate the minimal number of operations required for transforming one sequence into the other [62]. This defines an editing distance between gene sequences, which can be used to compare the evolution distance between species and construct the phylogenetic tree [61]. There have been many algorithms developed to determine genomic rearrangements. They consider different scenarios: whether the gene sequence is linear or circular, whether genes have unique labels, and what operations can be taken. Kececioglu and Sankoff only consider inversion for linear sequences with unique gene labels [34]; Blanchette et al. consider inversion and transposition for circular sequences with unique gene labels [6]; Tesler considers inversion, transposition, fusion, and fission for linear and circular sequences with unique gene labels [62]; Terauds and Sumner study circular sequences with representation theory tools [61]; Bohnenkämper et al. consider linear and circular sequences with possibly duplicated labels [8]. There are also systematic pipelines for determining rearrangements from whole-genome assemblies [19, 43]. Nevertheless, these methods consider large-scale rearrangements, and minimize the number of operations to transform one gene sequence into the other, not concrete genes that can change their locations. Besides, these methods only compare two gene sequences, not more. Their results depend on the set of possible operations, which is somewhat arbitrary.

In this paper, we consider a mesoscopic scenario between the genomic rearrangement situation and the short transposon situation: *Given accurately annotated gene sequences (not nucleotide sequences) from different individuals, determine individual genes (not short nucleotide segments or long gene strands) that can change their locations (transposable).* This provides a qualitative description for the stability of genes, which can guide gene editing [67] and phylogenetics [32]. The proportion of fixed genes quantifies the robustness of the genome. We aim at minimizing the number of genes to move. When there are only two gene sequences, this is equivalent to calculating genomic arrangements, where the only allowed operation is single-gene transposition.

In the copy-paste (duplication) case and deletion case, we can compare the numbers of copies of genes for different individuals to determine the transposable genes that have changed their copy numbers. In the inversion case, we can check the direction of genes to determine transposable genes that have changed their orientations [38]. In the cut-paste (insertion) case, the compositions of gene sequences are the same, but the orders of genes differ. It is not straightforward to uniquely determine which genes have changed their relative locations. Instead, we can consider the complement of transposable genes, which keep their relative locations and form a common subsequence of gene sequences from different individuals. Notice that genes in a subsequence does not need to be adjacent in the original sequences, different from a substring. We aim at explaining the difference among gene sequences with minimal transposable genes, meaning that we want to maximize the length of the complement of transposable genes. Thus we define the transposable genes to be the complement of the longest common subsequence. Given raw nucleotide sequences, we first transform them into gene sequences. Then we apply our algorithms to find the longest common subsequence, and the complement is transposable genes. If the longest common subsequence is not unique, we also need to determine which genes are more conserved and appear in all longest common subsequences.

It is common to use the length of the longest common subsequence as a quantitative score for comparing DNA sequences [12, 26, 83]. The longest common subsequence has also been used to define ultraconserved elements [55] or remove incongruent markers [16].

Determining the longest common subsequence is a classical problem in computer science. Various scenarios for this problem have been studied. Here we list Scenarios A-E, where the first two are more commonly studied. For more works in these scenarios, readers may refer to more thorough reviews [5, 24, 76]. Scenario A considers two sequences with possibly repeated genes, and the sequence length is *n*. The goal is to find the longest common subsequence, where the length is count by gene copies. This can be solved by dynamic programming with 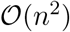 time complexity and 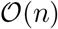 space complexity [23], but 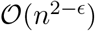 time complexity for any *ϵ* > 0 is impossible [4]. This also can be solved with *o*(*n*) space complexity and 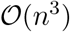 time complexity [35]. In Scenario *B*, there are *m* sequences with possibly repeated genes, and the sequence length is *n*. The goal is to find the longest common subsequence, where the length is count by gene copies. A standard dynamic programming algorithm has 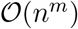 time complexity [7]. There have been other faster algorithms [66, 44, 27]. This scenario is equivalent to the maximum clique problem in graph theory, which is NP-hard [39], but has fast exact and heuristic algorithms [30, 37, 71]. Scenario C considers 2 sequences with possibly repeated genes, and the sequence length is n. The goal is to find the longest common subsequence, where each gene appears at most once. This scenario is NP-hard [1]. Scenario D is similar to Scenario B, but only consider common subsequences that contain or do not contain certain strings [68, 46]. In Scenario E, the gene sequences are arc-annotated, and the longest common subsequence should have the same arc annotation in original sequences [31].

In this paper, we consider four scenarios that are different from the previously studied longest common subsequence problems. These four scenarios are determined by two factors: whether the considered species has linear or circular gene sequences, and whether genes have multiple copies. When genes have multiple copies, we only consider common subsequences that consist of all or none of copies of the same gene. Scenario 1 has linear sequences without duplicated genes; Scenario 2 has circular sequences without duplicated genes; Scenario 3 has linear sequences with duplicated genes; Scenario 4 has circular sequences with duplicated genes.

Most known methods only aim at finding one longest common subsequence. When the longest common subsequence is not unique, we also need to classify whether a gene appears in all/some/none of the longest common subsequences. Determining all longest common subsequences is too time-consuming. To determine the relationship between genes and longest common subsequences, we develop corresponding algorithms with polynomial time complexities for Scenarios 1,2 (Algorithms 2,4). To our knowledge, there are no other determinations of whether genes appear in all longest common subsequences with polynomial complexities. Scenarios 3,4 only consider subsequences that consist of all or none copies of the same gene, and calculate the length by genes. Therefore, they are different from the classic Scenario B. We develop the equivalence of Scenario 3 with the maximum clique problems on graphs (Proposition 1). We prove that Scenario 4 is between the maximum clique problems on graphs and the maximum clique problems on 3-uniform hypergraphs (Propositions 2, 3). Although circular sequences are commonly studied in the context of genomic rearrangements, they are rare in the literature of longest common subsequence problems. Therefore, our Algorithm 3 that finds a longest common subsequence for Scenario 2 should also be novel. We test Algorithms 1,2,3,4 on the gene sequences of different Escherichia coli individuals and find some possible transposable genes.

If we only need to find one longest common subsequence, then Scenario 1 is a special case of Scenario B, and our method (Algorithm 1) is easily derived from standard algorithms. Scenarios 3,4 are equivalent to maximum clique problems in graphs and hypergraphs, which are NP-hard. These properties are also similar to Scenario B. Although there have been numerous algorithms for the maximum clique problem [77], for the sake of completeness, we design fast heuristic algorithms (Algorithms 5,6) and test them to find that they only fail in rare cases.

We proposed the idea of using the longest common subsequence to find transposable genes and Algorithm 1 in a previous paper [32], where Algorithm 1 was applied to study the “core-gene-defined genome organizational framework” (the complement of transposable genes) in various bacteria, and found that for different species, the transposable gene distribution and developmental traits are correlated. This paper considers other situations (especially when the longest common subsequence is not unique), and can be regarded as a theoretical sequel of that previous paper. Algorithm 1 is contained in this paper for the sake of completeness.

In sum, our main contributions are Algorithms 2,3,4 in Scenarios 1,2 and Propositions 1, 2, 3 in Scenarios 3,4.

We first describe the setup for the problem of determining transposable genes and transform it into the problem of finding the longest common subsequence. In the following four sections, we transform them into corresponding graph theory problems and design algorithms. We finish with some discussions. All the algorithms in this paper have been implemented in Python. For the code and data files, see https://github.com/YueWangMathbio/Transposon.

## 2 Setup

Given raw DNA sequencing data, the first step is to transform them into gene sequences. This can be done with various genome annotation tools [59, 9]. For simplicity, we replace the gene names by numbers 1,…,*n*.

For some species, the DNA is a line [58]. We can represent this DNA as a linear gene sequence of distinct numbers that represent genes: (1, 2, 3, 4). If some genes change their transcriptional orientations, we can simply detect them and handle the remaining genes. Now a linear DNA naturally has a direction (from 5’ end to 3’ end), thus (1, 2, 3, 4) and (4, 3, 2, 1) are two different gene sequences.

Consider two linear gene sequences from different individuals: (1, 2, 3, 4) and (1, 4, 2, 3). We can intuitively detect that gene 4 changes its relative position, and should be regarded as a transposable gene. However, changing the positions of genes 2, 3 can also transform one sequence into the other. The reason that we think gene 4 (not genes 2, 3) changes its relative position is that the number of genes we need to move is smaller. However, the number of genes that change their relative locations is difficult to determine. We can consider the complement of transposable genes, i.e., genes that do not change their relative positions. These fixed genes can be easily defined as the longest common subsequence of given gene sequences. Here a common subsequence consists of some genes (not necessarily adjacent, different from a substring) that keep their relative orders in the original sequences. *Thus transposable genes are the complement of this longest common subsequence.* Notice that the longest common subsequence might not be unique. We classify genes by their relations with the longest common subsequence(s). The motivation of classifying transposable genes with respect to the intersection and union of longest common subsequences is similar to defining essential variables with Markov boundaries in causal inference [73].

#### Definition 1.

*A gene is **proper-transposable** if it is not contained in any longest common subsequence. A gene is **non-transposable** if it is contained in every longest common subsequence. A gene is **quasi-transposable** if it is contained in some but not all longest common subsequences.*

In the example of (1, 2, 3, 4) and (1, 4, 2, 3), the unique longest common subsequence is (1, 2, 3). Thus 4 is proper-transposable, and 1, 2, 3 are non-transposable. In the following, we consider other scenarios, where the proper/quasi/non-transposable genes still follow Definition 1, but the definition of the longest common subsequence differs.

For some species, the DNA is a circle, not a line [65]. A circular DNA also has a natural direction (from 5’ end to 3’ end), and we use the clock-wise direction to represent this natural direction. In the circular sequence scenario, a common subsequence is a circular sequence that can be obtained from each circular gene sequence by deleting some genes. See Fig. 1 for two circular gene sequences and their longest common subsequence. Notice that we can rotate each circular sequence for a better match.

**Figure 1:**
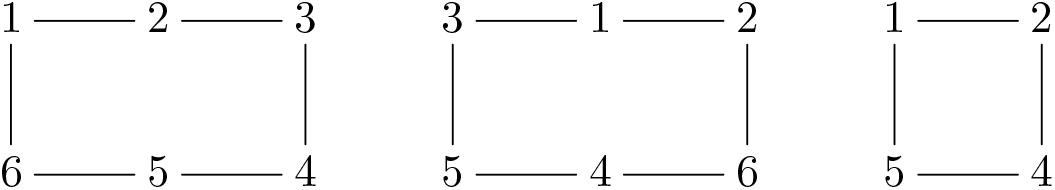
Two circular gene sequences without duplicated genes and their longest common subsequence, corresponding to Scenario 2.

A gene might have multiple copies (duplicated) in a gene sequence [25]. Notice that the definition of the transposable gene is a gene (specific DNA sequence) that has the ability to change its position, not a certain copy of a gene that changes its position. This means transposable genes should be defined for genes, not gene copies. Thus we should only consider common subsequences that consist of all or none copies of the same gene. When calculating the length of a common subsequence, we should count genes, not gene copies. Consider two linear sequences (4, 1, 2, 1, 1, 3, 2, 4, 1, 1) and (4, 1, 2, 3, 1, 1, 2, 1, 1, 4). If we consider any subsequences, the longest common subsequence is (4, 1, 2, 1, 1, 2, 1, 1); if we only consider subsequences that contain all or none copies of the same gene, but count the length by copies, the longest common subsequence is (1, 2, 1, 1, 2, 1, 1); if we only consider subsequences that contain all or none copies of the same gene, and count the length by genes, the unique longest common subsequence is (4, 2, 3, 2, 4), and gene 1 is proper-transposable.

When we consider circular gene sequences with duplicated genes, we should still only consider subsequences that consist of all or none copies of the same gene, and calculate the length by genes. Notice that circular sequences can be rotated. See Fig. 2 for two circular gene sequences with duplicated genes and their longest common subsequence.

**Figure 2:**
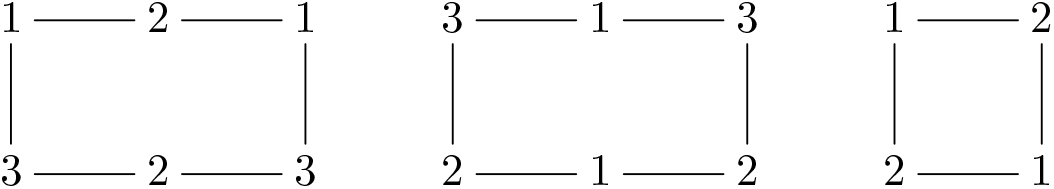
Two circular gene sequences with duplicated genes and their longest common subsequence, corresponding to Scenario 4.

We have turned the problem of determining transposable genes into finding the longest common subsequence of several gene sequences. Depending on whether the gene sequences are linear or circular, and whether genes have multiple copies, the problem can be classified into four scenarios:

**Scenario 1**: Consider *m* linear sequences of genes 1,…, *n*, where each gene has only one copy in each sequence. Determine the longest linear sequence that is a common subsequence of these *m* sequences.
**Scenario 2**: Consider *m* circular sequences of genes 1,…,*n*, where each gene has only one copy in each sequence. Determine the longest circular sequence that is a common subsequence of these *m* sequences. Here circular sequences can be rotated.
**Scenario 3**: Consider *m* linear sequences of genes 1,…,*n*, where each gene can have multiple copies in each sequence. Determine the longest linear sequence that is a common subsequence of these *m* sequences. Only consider subsequences that consist of all or none copies of the same gene, and calculate the length by genes.
**Scenario 4**: Consider *m* circular sequences of genes 1,… *,n,* where each gene can have multiple copies in each sequence. Determine the longest circular sequence that is a common subsequence of these *m* sequences. Only consider subsequences that consist of all or none copies of the same gene, and calculate the length by genes. Here circular sequences can be rotated.

These four scenarios correspond to different algorithms, and will be discussed separately.

## 3 Linear sequences without duplicated genes

In Scenario 1, consider *m* linear gene sequences, where each sequence contains *n* genes 1,…, *n*. Each gene has only one copy. For such permutations of 1,…, *n*, we need to find the longest common subsequence.

### 3.1 A graph representation of the problem

Brute-force searching that tests whether each subsequence appears in all sequences is not applicable, since the time complexity is exponential in *n*. To develop a polynomial algorithm, we first design an auxiliary directed graph 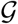.

#### Definition 2.

*For m linear sequences with n non-duplicated genes, the corresponding **auxiliary graph*** 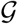 *is a directed graph, where each vertex is a gene g_i_, and there is a directed edge from g_i_ to g_j_ if and only if g_i_ appears before g_j_ in all m sequences.*

A directed path *g*_1_ → *g*_2_ → *g*_3_ → ⋯ → *g*_4_ → *g*_5_ in 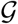 corresponds to a common subsequence (*g*_1_, *g*_2_, *g*_3_,…, *g*_4_, *g*_5_) of *m* sequences, and vice versa. We add 0 to the head of each sequence and *n* + 1 to the tail. Then the longest common subsequence must start at 0 and end at *n* + 1. *The problem of finding the longest common subsequence becomes finding the longest path from* 0 to *n* + 1 *in* 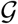 See Fig. 3 for an example of using the auxiliary graph to determine transposable genes. This auxiliary graph 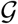 has no directed loop (acyclic). If there exists a loop *g*_1_ → *g*_2_ → *g*_3_ → ⋯ → *g*_4_ → *g*_1_, then *g*_1_ is prior to *g*_4_ and *g*_4_ is prior to *g*_1_ in all sequences, a contradiction.

**Figure 3:**
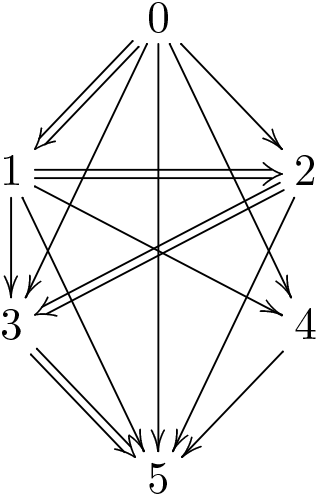
The auxiliary graph 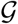 of two sequences ([0], 1, 2, 3, 4, [5]) and ([0], 1, 4, 2, 3, [5]). The unique longest path (double arrows) from 0 to 5 is 0 → 1 → 2 → 3 → 5, meaning that the unique longest common sequence is ([0], 1, 2, 3, [5]). Thus 1, 2, 3 are non-transposable, and 4 is proper-transposable.

### 3.2 Find the longest path

Determining the longest path between two vertices in a directed acyclic graph can be solved by a standard dynamic programming algorithm. For a vertex *g_i_* ∈ {0,1,…,*n*}, consider the longest path from *g_i_* to *n* + 1. Since there exists an edge *g_i_* → *n* + 1, and 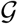 is acyclic, this longest path exists. If the longest path is not unique, assign one arbitrarily.

#### Definition 3.

*Define F*_+_(*g_i_*) *to be the length of the longest path from g_i_ to n* + 1 *in* 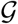, *and H*_+_(*g_i_*) *to be the vertex next to g_i_ in this path*.

*F*_+_and *H*_+_ can be calculated recursively: For one gene *g_i_*, consider all genes *g_j_* with an edge *g_i_* → *g_j_* in 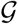. The gene *g_j_* with the largest *F*_+_(*g_j_*) is assigned to be *H*_+_(*g_i_*), and *F*_+_(*g_i_*) = *F*_+_(*g_j_*) + 1. If *g_l_* → *n* + 1 is the only edge that starts from gene *g_l_*, then *F*_+_(*g_l_*) = 1, and *H*_+_(*g_l_*) = *n* + 1. In other words,

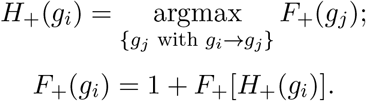

Then 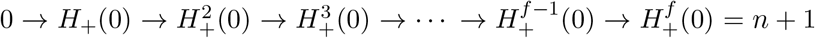, denoted by 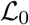, is a longest path in 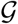. Here *f* = *F*_+_(0), and 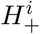 is the *i*th iteration of *H*_+_.

### 3.3 Test the uniqueness of the longest path

To test whether quasi-transposable genes exist, we need to check the uniqueness of this longest path.

#### Definition 4.

*For g_i_* ∈ {1,…,*n, n* + 1}, *define F*___(*g_i_*) *to be the length of the longest path from* 0 *to g_i_ in* 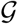, *and H*___(*g_i_*) *to be the vertex prior to g_i_ in this path*.

*F*___ and *H*___ can be calculated similar to *F*_+_ and H_+_. We can see that *F*_+_(*g_i_*) + *F*___(*g_i_*) is the length of

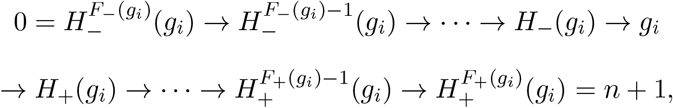

a longest path from 0 through *g_i_* to *n* + 1. For 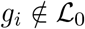, if *F*_+_(*g_i_*) + *F*___(*g_i_*) < *F*_+_(0), then *g_i_* is proper-transposable; if *F*_+_(*g_i_*) + *F*___(*g_i_*) = *F*_+_(0), then *g_i_* is quasi-transposable. If every 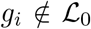 is proper-transposable, then the longest common subsequence is unique, and all genes in 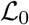 (excluding the auxiliary 0 and *n* + 1) are non-transposable. The procedure of determining transposable genes stops here. Otherwise, the longest common subsequence is not unique, and we need to find quasi-transposable genes in 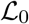.

### 3.4 Find quasi-transposable genes

When determining all quasi-transposable genes *g*_1_,…,*g_k_* not in 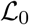, as described above, we construct corresponding longest paths 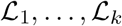 from 0 to *n* + 1, where each 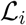 passes through *g_i_*. We claim that a gene 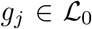 is non-transposable if and only if *g_j_* is contained in all 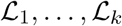. To prove this, we need the following lemma.

#### Lemma 1.

*In Scenario 1 of linear sequences without duplicated genes, each quasi-transposable gene g_i_ has a corresponding quasi-transposable gene g_j_, so that no longest common subsequence can contain both g_i_ and g_j_*.

If a gene 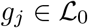 is non-transposable, then it is contained in all 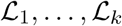. If 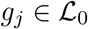 is quasi-transposable, by Lemma 1, there is a quasi-transposable gene 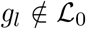 which is mutual-exclusive with *g_j_*, in the sense that *g_l_* and *g_j_* cannot appear in the same longest common subsequence. The corresponding longest path 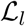 contains *g_l_*, thus cannot contain *g_j_*. This proves our approach to determine the quasi-transposable genes in 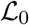.

#### Proof of Lemma 1.

Fix a quasi-transposable gene *g_i_*. It is contained in a longest path 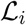, which contains all non-transposable genes. Thus for each non-transposable gene *g*\*, there is an edge between *g*\* and *g_i_* in 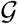. Assume *g_i_* has no such mutual-exclusive quasi-transposable gene *g_j_*. Then there is an edge (direction unknown) in 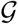 between *g_i_* and each quasi-transposable gene *g_j_*. Choose a longest path 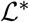 in 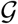 that does not contain *g_i_*. Whether 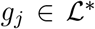 is a non-transposable gene or a quasi-transposable gene, there is an edge between *g_j_* and *g_i_*. Determine the first gene *g_k_* in 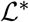 that has an edge *g_i_* → *g_k_*. Since there is an edge *g_i_* → *n* + 1, *g_k_* exists. Since there is an edge 0 → *g_i_*, *g_k_* = 0. Denote the previous gene of *g_k_* in 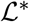 by *g_l_*, then *g_l_* exists, and there is an edge *g_l_* → *g_i_* Thus we construct a path 0 →⋯→ *g_l_* → *g_i_* → *g_k_* →⋯→ *n* + 1, which is longer than the longest path, a contradiction. Thus *g_i_* has a mutual-exclusive quasi-transposable gene *g_j_*.

### 3.5 Algorithms and complexities

We summarize the above method as Algorithms 1,2. If we have known that the longest common subsequence is unique, then we just need to apply Algorithm 1, so that genes in 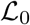 are non-transposable, and genes not in 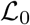 are proper-transposable. We have reported Algorithm 1 previously [32, 69]. Algorithm 1 is kept here to make the story complete. Assume we have *m* sequences with length *n*, and the length of the longest common subsequence is *n* – *k*. The time complexities of Steps 2-5 in Algorithm 1 are 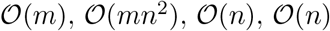. The time complexities of Step 2 and Step 3 in Algorithm 2 are 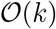 and 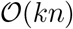. Since *k* ≤ *n*, the overall time complexity of determining transposable genes in Scenario 1 by Algorithms 1,2 is 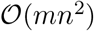. The space complexity is trivially 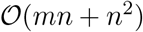.

### 3.6 Applications on experimental data

We test Algorithms 1,2 on *Escherichia coli* gene sequences. From NCBI sequencing database, we obtain gene sequences of three individuals of *E. coli* strain ST540 (GenBank CP007265.1, GenBank CP007390.1, GenBank CP007391.1) and three individuals of *E. coli* strain ST2747 (GenBank CP007392.1, GenBank CP007393.1, GenBank CP007394.1).

All three sequences of ST540 start with gene dnaA and end with gene rpmH. We can regard them as linear gene sequences. We remove genes that appear more than once in one sequence, and remove genes that do not appear in all three sequences. After applying Algorithms 1,2 on these three sequences, there are 301 non-transposable genes, 4 quasi-transposable genes (hpaC, iraD, fbpC, psiB), and 263 proper-transposable genes. The reason for the large amount of proper-transposable genes is that sequence CP007265.1 is significantly different from the other two. After removing it and applying Algorithms 1,2 to the remaining two sequences (CP007390.1 and CP007391.1), there are 564 non-transposable genes and 4 quasi-transposable genes (hpaC, iraD, fbpC, psiB). Therefore, some genes in hpaC, iraD, fbpC, psiB are likely to translocate.

**Algorithm 1:**
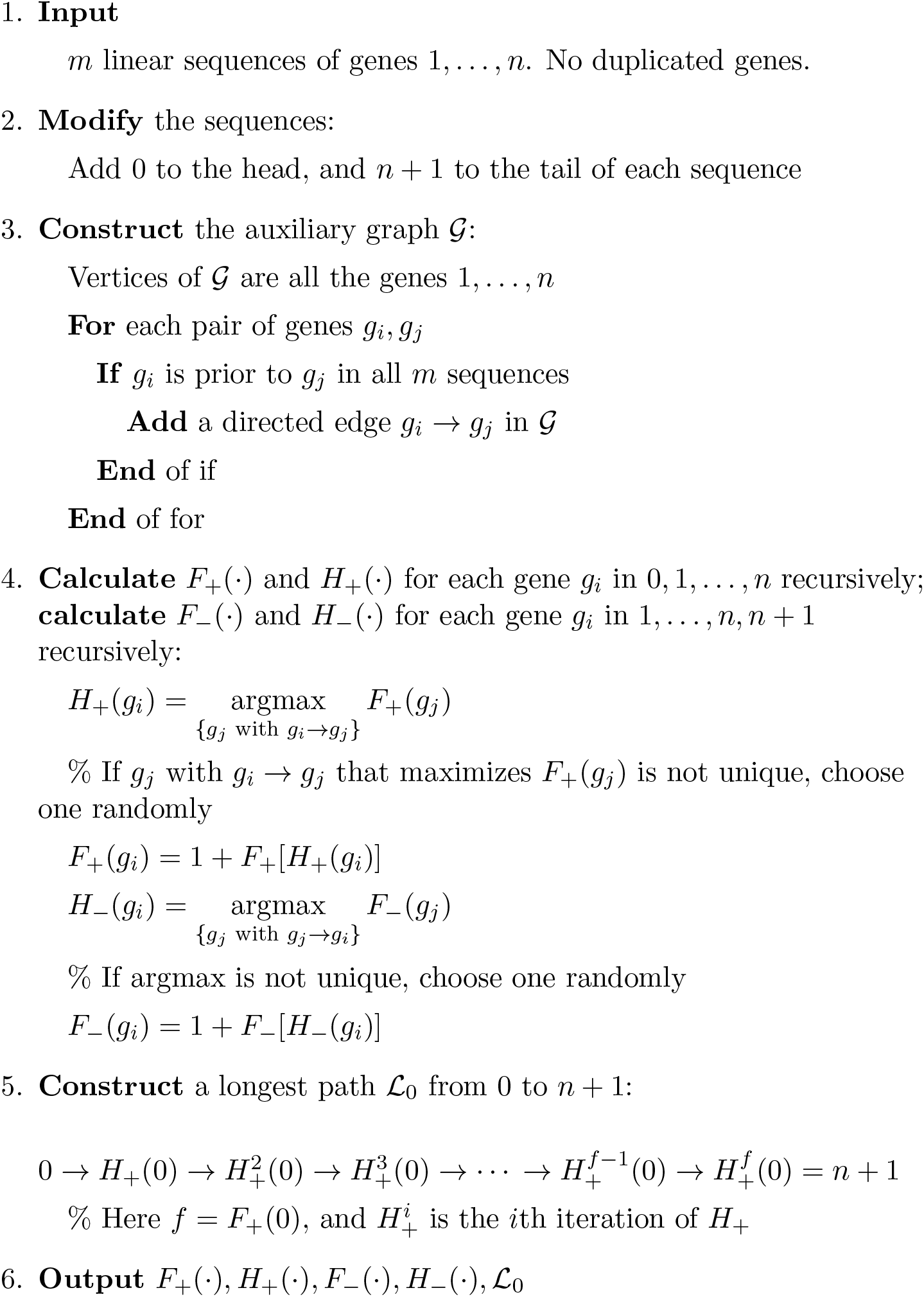
Detailed workflow of determining proper-transposable genes and quasi-transposable genes in Scenario 1, preparation stage.

**Algorithm 2:**
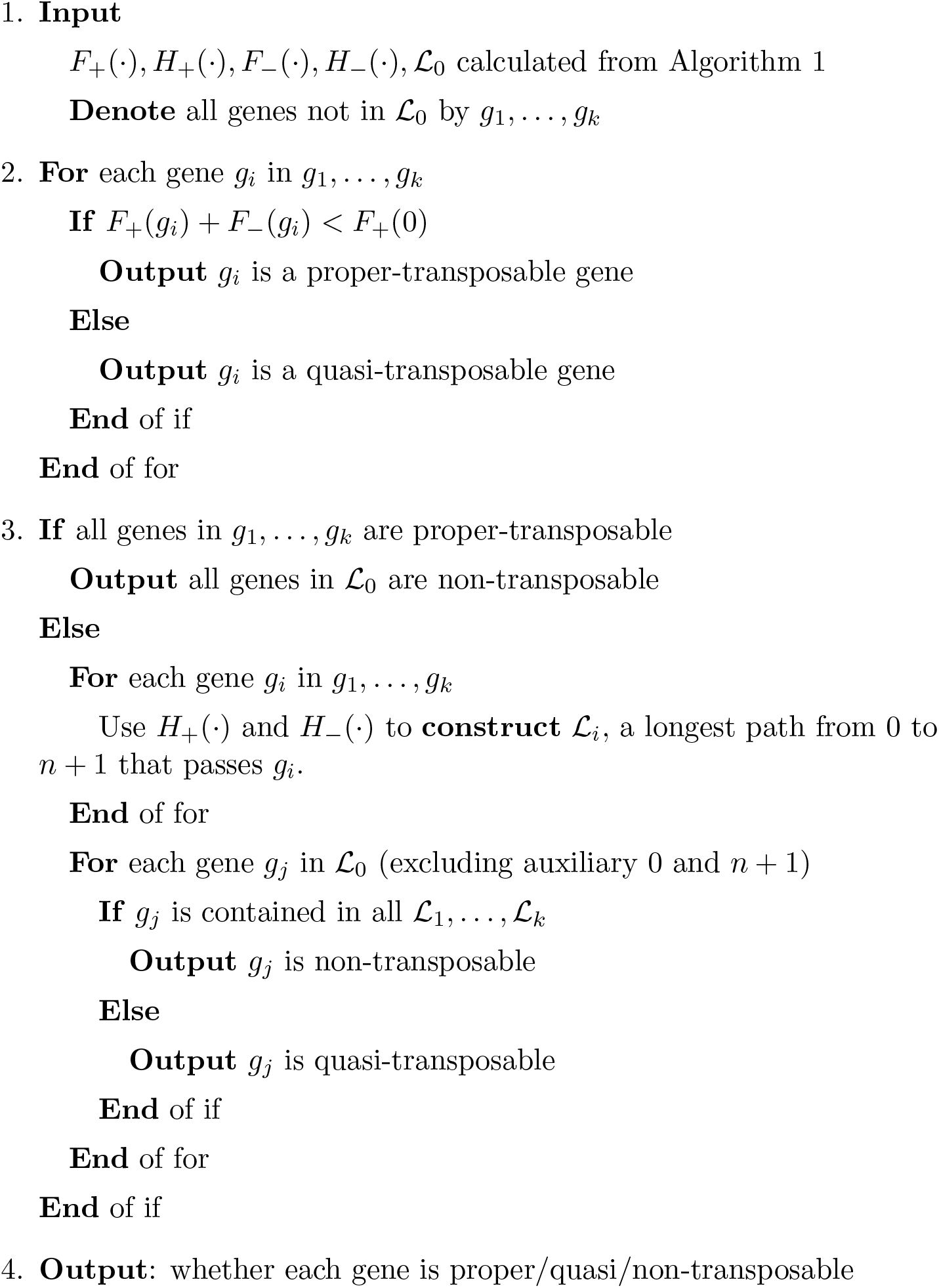
Detailed workflow of determining proper-transposable genes and quasi-transposable genes in Scenario 1, output stage.

All three sequences of ST2747 start with gene glnG and end with gene hemG. We can regard them as linear gene sequences. We remove genes that appear more than once in one sequence, and remove genes that do not appear in all three sequences. After applying Algorithms 1,2 on these three sequences, all 573 genes are non-transposable.

## 4 Circular sequences without duplicated genes

In Scenario 2, consider *m* circular gene sequences, where each sequence contains *n* genes 1,…, *n*. Each gene has only one copy in each sequence. For such circular permutations of 1,…, *n*, we need to find the longest common subsequence. Assume the length of the longest common subsequence is *n*–*k*.

### 4.1 Find a longest common subsequence

We first randomly choose a gene *g_i_*. Cut all circular sequences at *g_i_* and expand them to be linear sequences. For example, the circular sequences in Fig. 1 cut at 1 are correspondingly (1, 2, 3, 4, 5, 6) and (1, 2, 6, 4, 5, 3). Using Algorithm 1, we can find 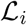that begins with *g_i_*, which is a longest common subsequence of all expanded linear sequences. In the above example, the longest common linear subsequence starting from 1 is (1, 2, 4, 5). If *g_i_* is a non-transposable gene or a quasi-transposable gene, then 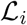 (glued back to a circle) is a longest common circular subsequence. If *g_i_* is a proper-transposable gene, then 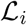 is shorter than the longest common circular sub-sequence. In Fig. 1, gene 1 is non-transposable, and (1, 2, 4, 5) (glued) is the longest common circular subsequence.

We do not know if 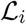 (glued) is a longest common subsequence (whether containing *g_i_* or not) for all circular sequences. If there is a longer common subsequence, it should contain genes that are not in 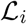. Consider four variables 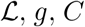, and 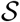, whose initial values are 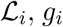, *g_i_*, the length of 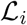, and the complement of 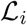. These variables contain information on the longest common linear subsequence that we have found during this procedure.

Choose a gene *g_j_* in 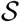, and cut all circular gene sequences at *g_j_*. Apply Algorithm 1 to find 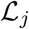, which is the longest in common subsequences that contain *g_j_*. If the length of 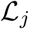 is larger than *C*, set 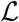 to be 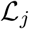, set *g* to be *g_j_*, set *C* to be the length of 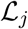, and set 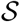 to be the complement of 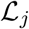. Otherwise, keep 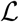, *g, C*, and 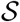 still.

Choose another gene *g_l_* in 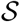 which has not been chosen before, and repeat this procedure. This procedure terminates when all genes in 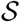 have been chosen and cut. Denote the final values of 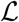, *g, C*, and 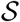 by 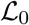, *g*_0_, *C*_0_, and 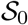. Here 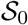 is the complement of 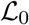.

During this procedure, if the current *g* is a proper-transposable gene, then 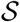 contains a non-transposable gene or a quasi-transposable gene, which has not been chosen. Thus 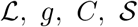 will be further updated. If the current *g* is a non-transposable gene or a quasi-transposable gene, then *C* has reached its maximum, and 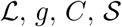 will not be further updated. This means 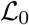 is a longest common circular subsequence, and *C*_0_ is the length of the longest common subsequence, *n* – *k*. Also, the total number of genes being chosen and cut is *k* + 1. All *k* genes in 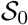 and *g*_0_ are chosen and cut. A gene *g_t_* in 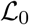 (excluding *g*_0_) is non-transposable or quasi-transposable, and cannot be chosen and cut. The reason is that it cannot be chosen before *g*_0_ is chosen (only proper-transposable genes can be chosen before *g*_0_ is chosen), and it cannot be chosen after *g*_0_ is chosen 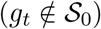.

### 4.2 Determine quasi-transposable genes

For each gene 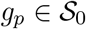, apply Algorithm 1 to calculate *C_p_*, the length of the longest common subsequence that contains *g_p_*. If *C_p_* < *C*_0_, *g_p_* is a proper-transposable gene. Otherwise, *C_p_* = *C*_0_ means *g_p_* is a quasi-transposable gene. We have found all proper-transposable genes. If all genes in 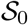 are proper-transposable, then all genes in 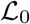 are non-transposable, and the procedure terminates.

If 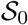 contains quasi-transposable genes, then 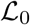 also has quasi-transposable genes. To determine quasi-transposable genes in 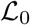, we need the following lemma.

#### Lemma 2.

*In Scenario 2, choose a quasi-transposable gene g_p_ and cut the circular sequences at g_p_ to obtain linear sequences. A proper-transposable gene for the circular sequences is also a proper-transposable gene for the linear sequences; a non-transposable gene for the circular sequences is also a non-transposable gene for the linear sequences.*

*Proof.* Consider a longest common subsequence 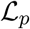 for linear sequences cut at *g_p_*. Since *g_p_* is a quasi-transposable gene, the length of 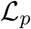 is also *n* – *k*, meaning that 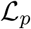 is also a longest common subsequence for circular sequences. Now, this lemma is proved by the definition of proper/quasi/non-transposable gene.

If a gene *g_r_* in 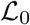 is non-transposable for the circular sequences, then *g_r_* is a non-transposable gene for linear sequences cut at each quasi-transposable gene 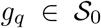. If a gene *g_s_* in 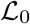 is quasi-transposable for the circular sequences, then there is a longest common circular subsequence 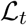 that does not contain *g_s_*, meaning that 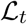 contains a quasi-transposable gene *g_t_* not in 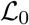. Then *g_s_* is a proper/quasi-transposable gene for linear sequences cut at *g_t_*.

Therefore, we can use the following method to determine quasi-transposable genes in 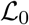. For each quasi-transposable gene 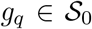, cut at *g_q_* and apply Algorithms 1,2 to determine if each gene in 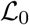 is proper/quasi/non-transposable for the linear gene sequences cut at *g_q_*. A gene 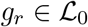 is non-transposable for the circular sequences if and only if it is non-transposable for linear sequences cut at any quasi-transposable gene 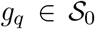. A gene 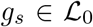 is quasi-transposable for the circular sequences if and only if it is proper/quasi-transposable for linear sequences cut at some quasi-transposable gene 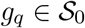.

When we have determined all quasi-transposable genes in 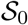, it might be tempting to apply a simpler approach to determine quasi-transposable genes in 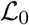: For each quasi-transposable gene 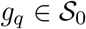, cut at *g_q_* and apply Algorithm 1 to find a longest common subsequence 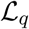. A gene in 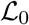 is non-transposable if and only if it appears in all such 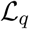. This approach is valid only if the following conjecture holds, which is similar to Lemma 1:

#### Conjecture 1.

*In Scenario 2 of circular sequences without duplicated genes, each quasi-transposable gene g_i_ has a corresponding quasi-transposable gene g_j_, so that no longest common subsequence can contain both g_i_ and g_j_*.

However, Conjecture 1 does not hold. See Fig. 4 for a counterexample. All genes are quasi-transposable. Any two quasi-transposable genes are contained in a longest common subsequence (length 3). Thus the simplified approach above does not work.

**Figure 4:**
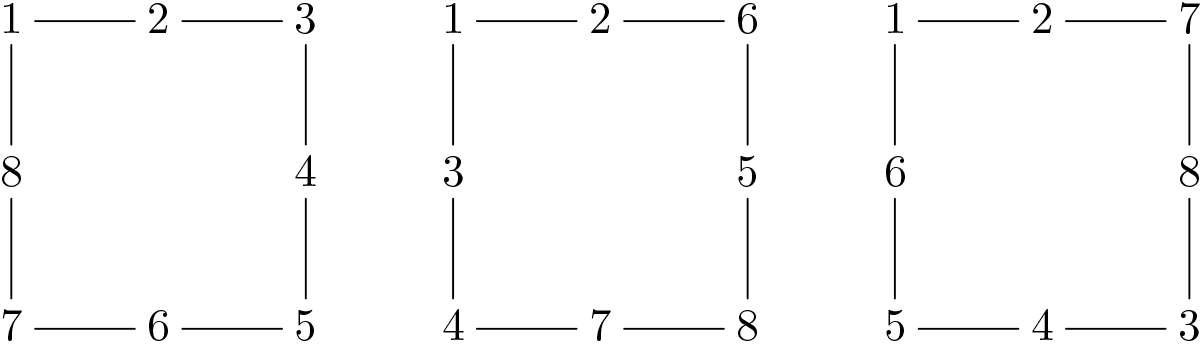
A counterexample with three circular sequences that fails Conjecture 1.

We summarize the above method as Algorithms 3,4. If we have known that the longest common subsequence is unique, then we just need to apply Algorithm 3, so that genes in 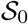 are proper-transposable, and genes not in 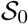 are non-transposable. Assume we have *m* sequences with length *n*, and the length of the longest common subsequence is *n* – *k*. The time complexities of Step 2 and Step 3 in Algorithm 3 are 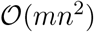 and 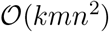. The time complexities of Step 2 in Algorithm 4 is 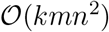. The overall time complexity of determining transposable genes in Scenario 2 by Algorithms 3,4 is 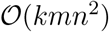. The space complexity is trivially 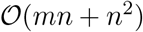.

### 4.3 Applications on experimental data

Similar to Subsection 3.6, we test Algorithms 3,4 on *Escherichia coli* gene sequences. From NCBI sequencing database, we obtain gene sequences of three individuals of *E. coli* strain ST540 (GenBank CP007265.1, GenBank CP007390.1, GenBank CP007391.1) and three individuals of *E. coli* strain ST2747 (GenBank CP007392.1, GenBank CP007393.1, GenBank CP007394.1).

We regard all three sequences of ST540 as circular gene sequences. We remove genes that appear more than once in one sequence, and remove genes that do not appear in all three sequences. After applying Algorithms 3,4 on these three sequences, there are 389 non-transposable genes, 50 quasi-transposable genes, and 129 proper-transposable genes. The reason for the large amount of proper-transposable genes is that sequence CP007265.1 is significantly different from the other two. After removing it and applying Algorithms 3,4 to the remaining two sequences (CP007390.1 and CP007391.1), there are 564 non-transposable genes and 4 quasi-transposable genes (hpaC, iraD, fbpC, psiB). Therefore, some genes in hpaC, iraD, fbpC, psiB are likely to translocate.

We regard all three sequences of ST2747 as circular gene sequences. We remove genes that appear more than once in one sequence, and remove genes that do not appear in all three sequences. After applying Algorithms 3,4 on these three sequences, all 573 genes are non-transposable genes.

**Algorithm 3:**
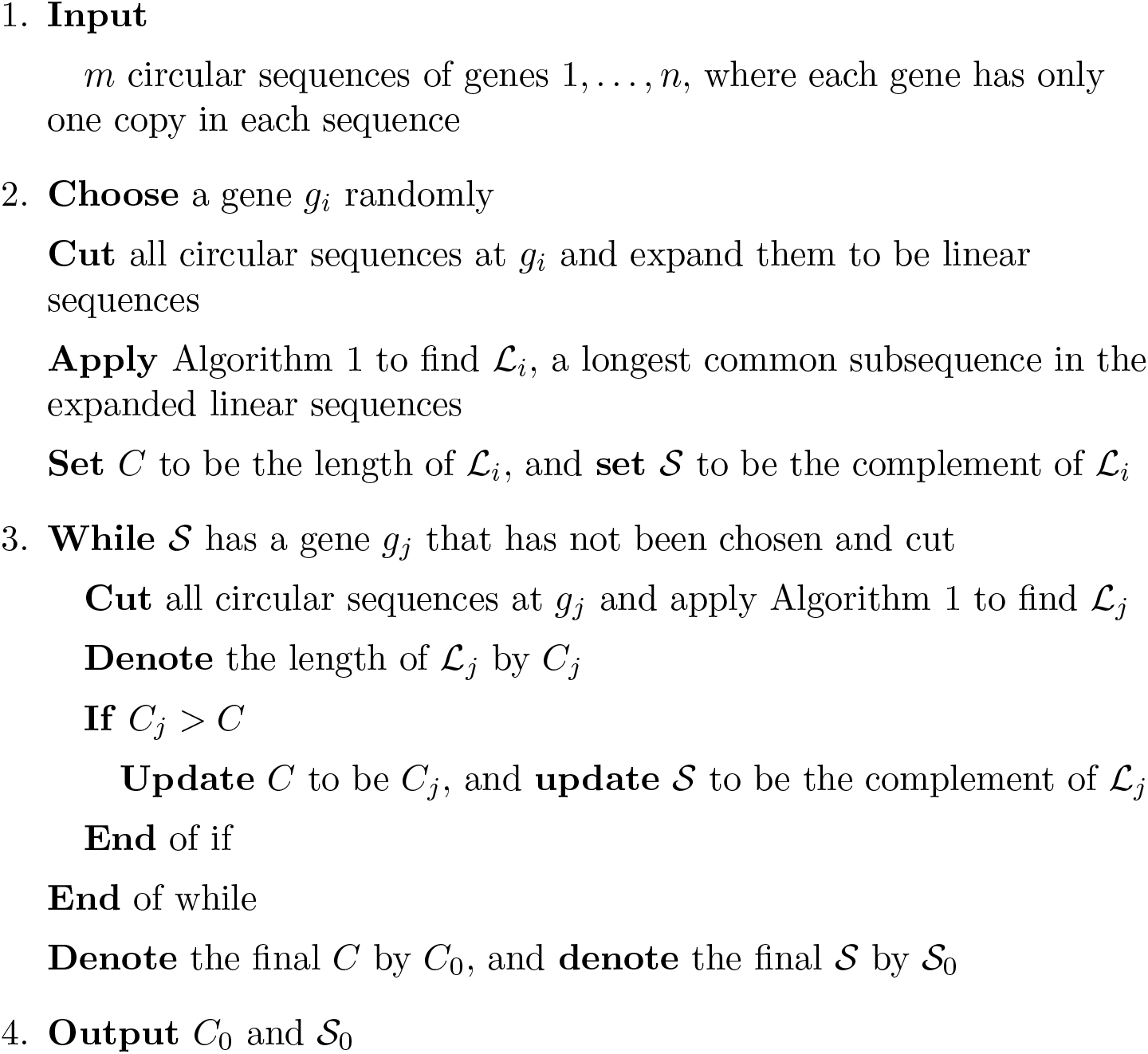
Detailed workflow of determining proper-transposable genes and quasi-transposable genes in Scenario 2, preparation stage.

**Algorithm 4:**
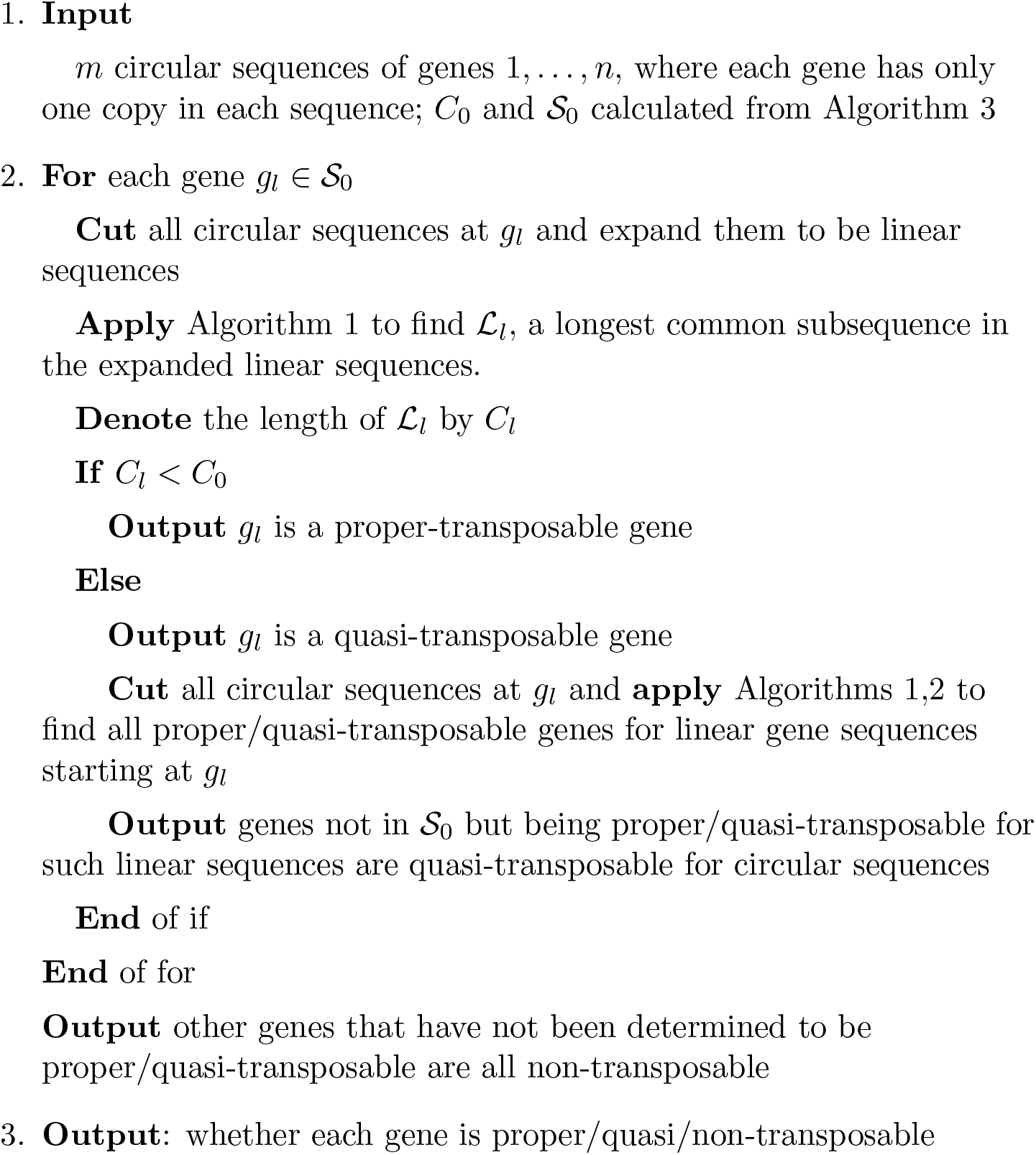
Detailed workflow of determining proper-transposable genes and quasi-transposable genes in Scenario 2, output stage.

## 5 Linear sequences with duplicated genes

In Scenario 3, consider *m* linear gene sequences, where each sequence contains different numbers of copies of *n* genes 1,…,*n*. We need to find the longest common subsequence. Here we only consider common subsequences that consist of all or none copies of the same gene, and the subsequence length is calculated by genes, not gene copies.

### 5.1 A graph representation of the problem

Similar to Scenario 1, we construct an auxiliary graph 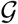, where each vertex is a gene (not a copy of a gene). However, in this case, the auxiliary graph is undirected: There is an undirected edge between gene *g_i_* and gene *g_j_* if and only if all the copies of *g_i_* and *g_j_* keep their relative locations in all sequences. For example, consider two sequences (1, 2, 3, 2, 3, 4, 5) and (2, 1, 3, 3, 2, 4, 5). For gene pair 1, 3, the corresponding sequences are (1, 3, 3) and (1, 3, 3), meaning that there is an edge between 1 and 3. For gene pair 1, 2, the corresponding sequences are (1, 2, 2) and (2, 1, 2), meaning that there is no edge between 1 and 2. See Fig. 5 for the auxiliary graph in this case.

**Figure 5:**
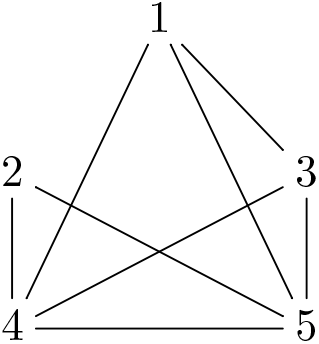
The auxiliary graph 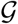 of two sequences (1, 2, 3, 2, 3, 4, 5) and (2, 1, 3, 3, 2, 4, 5). The unique largest complete subgraph is {1, 3, 4, 5}, meaning that the unique longest common sequence is (1, 3, 3, 4, 5). Thus 1, 3, 4, 5 are non-transposable genes, and 2 is a proper-transposable gene.

#### Definition 5.

*A subgraph of* 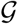 *consists of some genes g*_1_,…,*g_l_ and the edges between them. In a subgraph, if there is an edge between any two genes, this subgraph is called a complete subgraph (also called a clique)*.

#### Definition 6.

*In graph* 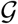, *the degree of a gene g is the number of edges linking g. In a complete graph of p genes, where any two genes have an edge in between, each gene has degree p – 1.*

#### Definition 7.

*If all copies of genes g_1_,…, *g_l_ keep their relative locations in all linear sequences, we say that g*_1_,…, g_l_ form a common subsequence*.

The following Lemma 3 shows that there is a bijection between common subsequences and complete subgraphs in 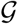. *The problem of determining the longest common subsequence now becomes determining the largest complete subgraph of* 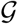.

#### Lemma 3.

*In Scenario 3, construct the auxiliary graph* 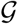 *from gene sequences. If g*_1_,…, *g_k_ form a complete subgraph in* 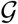, *then g*_1_,…, *g_k_ form a common subsequence, and vice versa*.

*Proof.* If *g*_1_,…,*g_l_* form a common subsequence, then there is an edge in 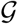 between any two genes in *g*_1_,…, *g_l_*, meaning that they form a complete subgraph.

For the other direction, only consider copies of *g*_1_,…,*g_k_* in these sequences. If *g*_1_,…,*g_k_* do not form a common subsequence, find the first digit that such sequences differ. Assume *g_p_* and *g_q_* can both appear in this digit. Then *g_p_, g_q_* cannot form a common subsequence, and there is no edge between *g_p_* and *g_q_*.

We illustrate this proof with Fig. 5: For genes 2, 3, 4, the sequences are (2, 3, 2, 3, 4) and (2, 3, 3, 2, 4). The third digit is different, where 2 and 3 can both appear. Then the sequences for genes 2, 3, (2, 3, 2, 3) and (2, 3, 3, 2), cannot match, and there is no edge between 2 and 3.

### 5.2 A heuristic algorithm

The above discussion shows that given gene sequences, we can construct an undirected graph 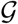, so that there is a bijection between common subsequences and complete subgraphs. The inverse also holds: We can construct corresponding gene sequences for a graph.

#### Lemma 4.

*Given an undirected graph* 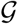, *we can construct two gene sequences, so that there is a bijection between common subsequences and complete subgraphs.*

*Proof.* Assume the graph has *n* genes. We start with two sequences (1, 2,…, *n*) and(1, 2,…,*n*). For each pair of genes *g*_i_,g_j_·, if there is no edge between them in 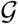, add *g_i_, g_j_* to the end of the first sequence, and *g_j_, g_i_* to the end of the second sequence. Then *g_i_, g_j_* cannot both appear in a common subsequence, and this operation does not affect other gene pairs.

For example, corresponding to Fig. 5, we start with (1, 2, 3, 4, 5) and (1, 2, 3, 4, 5). Since there is no edge between 1, 2, we add them to have (1, 2, 3, 4, 5, 1, 2) and (1, 2, 3, 4, 5, 2, 1). Since there is no edge between 2, 3, we add them to have (1, 2, 3, 4, 5, 1, 2, 2, 3) and (1, 2, 3, 4, 5, 2, 1, 3, 2). These two sequences corresponds to Fig. 5.

Combining Lemma 3 and Lemma 4, we obtain the following result:

#### Proposition 1.

*Finding the longest common sequence in Scenario 3 is equivalent to the maximum clique problem, which is NP-hard.*

*Proof.* For an undirected graph, we can use Lemma 4 to construct corresponding sequences. If we have the solution of finding the longest common sequence in Scenario 3, then we can find the largest complete subgraph in an extra polynomial time.

For gene sequences in Scenario 3, we can construct corresponding auxiliary graph. If we have the solution of finding the largest complete subgraph, then we can use Lemma 3 to find the longest common sequence in Scenario 3 in an extra polynomial time.

Therefore, finding the longest common sequence in Scenario 3 and finding the largest complete subgraph are equivalent. The problem of determining the largest complete subgraph is just the maximum clique problem, which is NP-hard [64]. Thus finding the longest common sequence in Scenario 3 is also NP-hard. This means it is not likely to design an algorithm that always correctly determines the longest common subsequence in polynomial time.

We have transformed Scenario 3 into the maximum clique problem for a graph 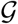. There have been various algorithms for the maximum clique problem [30, 37, 71], and readers may refer to a review for more details [77]. For completeness, we propose a simple idea: In the auxiliary graph 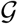, repeatedly abandon the gene with the smallest degree (and also edges linking this gene) until the remaining genes form a complete subgraph. See Algorithm 5 for the details of this greedy heuristic method. This algorithm is easy to understand, and can provide some intuition. We do not claim that Algorithm 5 is comparable to other sophisticated algorithms.

We test Algorithm 5 on random graphs. Construct a random graph with *n* genes, and any two genes have probability 0.5 to have an edge in between. Use brute-force search to find the maximum clique, and compare its size with the result of Algorithm 5. For each *n* ≤ 15, we repeat this for 10000 times, and every time Algorithm 5 returns the correct result. Therefore, for small random graphs, the 95% credible interval for the success rate of Algorithm 5 is [0.9997, 1]. We can claim that Algorithm 5 is a good heuristic algorithm that fails with a very small probability. Since finding the true maximum clique requires exponentially slow brute-force search, we do not test on very large graphs.

**Algorithm 5:**
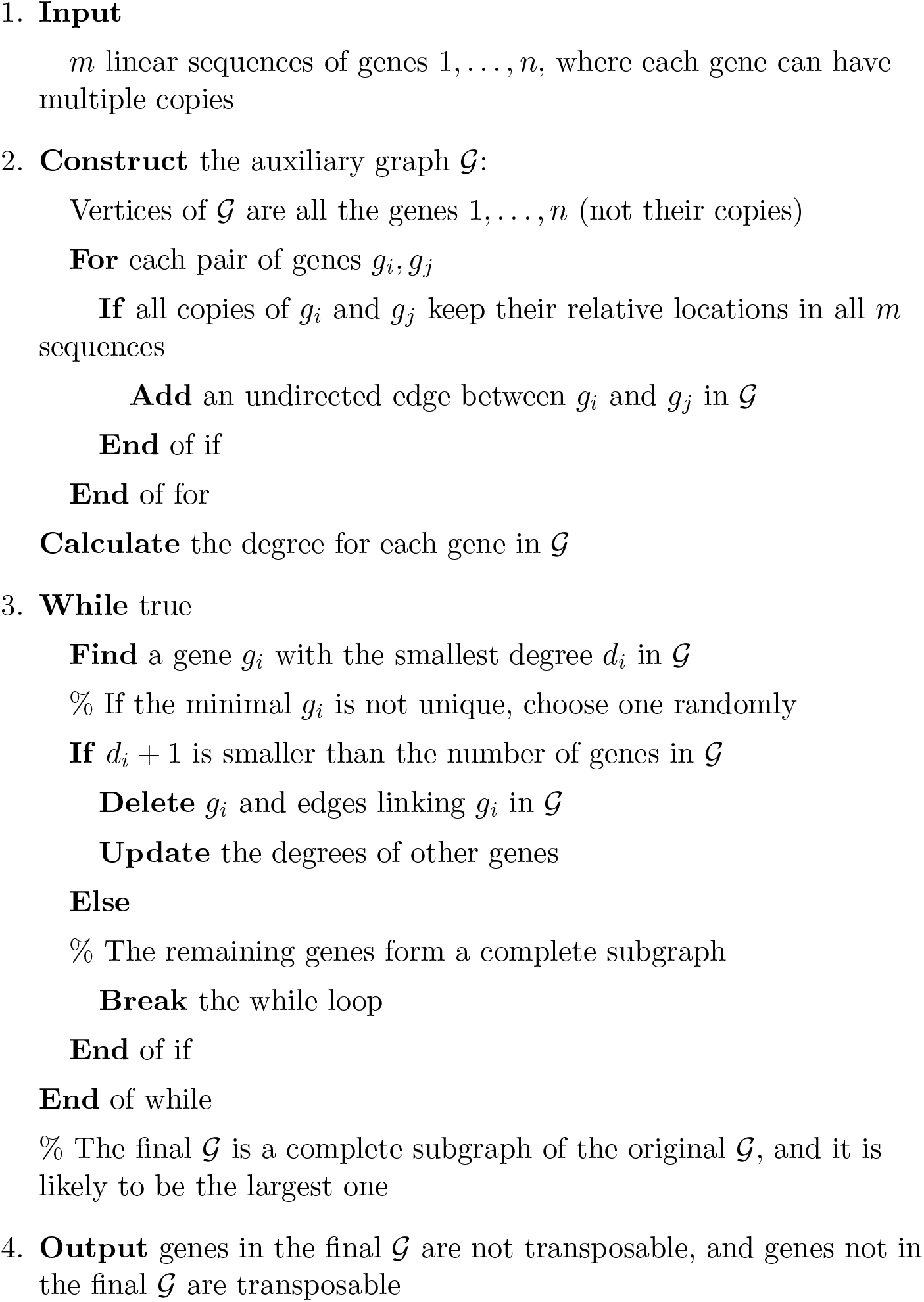
A heuristic method for detecting transposable genes in Scenario 3.

Nevertheless, Algorithm 5 does not always produce the correct result. See Fig. 6 for a counterexample. Here genes 1, 2, 3, 4, 5, 6 have degree 4, while genes 7, 8, 9, 10 have degree 3. When applying Algorithm 5, genes 7, 8, 9, 10 are first abandoned, and the final result just has three genes, such as 1, 3, 5. However, the largest complete graph is 7, 8, 9, 10. Besides, Algorithm 5 can only determine one (possibly longest) common subsequence. Thus we cannot determine the existence of quasi-transposable genes.

**Figure 6:**
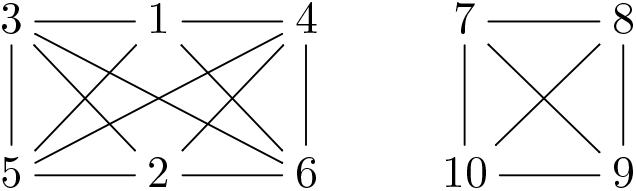
The auxiliary graph 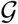 of linear sequences (7, 8, 9, 10, 1, 1, 2, 3, 3, 4, 5, 5, 6) and (1, 2, 1, 3, 4, 3, 5, 6, 5, 7, 8, 9, 10). This counterexample fails Algorithm 5.

Assume we have *m* sequences with *n* genes. In general, the copy number of a gene is small, and we can assume the length of each sequence is 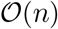. The time complexities of Step 2 and Step 3 in Algorithm 5 are 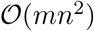 and 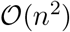, and the overall time complexity is 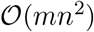. The space complexity is trivially 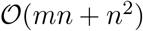.

## 6 Circular sequences with duplicated genes

In Scenario 4, consider *m* circular gene sequences, where each sequence contains different numbers of copies of *n* genes 1,…, *n*. We need to find the longest common subsequence. Here we only consider common subsequences that consist of all or none copies of the same gene, and the subsequence length is calculated by genes, not gene copies.

We shall prove that finding the longest common subsequence in Scenario 4 is no easier than in Scenario 3. Thus Scenario 4 is also NP-hard.

#### Proposition 2.

*Finding the longest common subsequence in Scenario 4 is NP-hard.*

*Proof.* From Proposition 1, Scenario 3 is NP-hard, meaning that any NP problem can be reduced to Scenario 3 in polynomial time. We just need to prove that Scenario 3 can be reduced to Scenario 4 in polynomial time.

Given *m* linear sequences with *n* genes in Scenario 3, add genes *n* + 1,…, 2*n* + 1 to the end of each sequence, and glue each linear sequence into a circular sequence. The longest common subsequence for these circular sequence has the following properties: (1) it contains all genes *n*+1,…, 2*n*+ 1; (2) after cutting at *n* + 1 and removing genes *n* + 1,…, 2*n* + 1, the remaining linear sequence is the longest common subsequence in Scenario 3.

(1) The longest common subsequence has at least *n* + 1 genes (*n* + 1,…, 2*n* + 1). Therefore, at least one gene in *n* + 1,…, 2*n* + 1 is included, such as *n* + 1. Since gene *n* + 1 aligned in all sequences, *n* + 2,…, 2*n* + 1 are also aligned, meaning that they are also in the longest common subsequence.

(2) After cutting and removing *n* + 1,…, 2*n* + 1, the remaining linear sequence is a common subsequence in Scenario 3. If there is a longer common subsequence, then that with *n* + 1,…, 2*n* + 1 should be a longer common subsequence in Scenario 4, a contradiction.

Therefore, if we can find the longest common subsequence for these circular sequences, then we can find the longest common subsequence for linear sequences in polynomial time.

Similar to Scenario 3, to find the longest common subsequence in Scenario 4, we want to reduce it to a maximum clique problem. However, Lemma 3 does not hold in Scenario 4. For example, we can consider a circular sequence (1, 2, 3) and its mirror symmetry. These two sequences are different, but any two genes form a common subsequence. However, inspired by Lemma 3, we have the following conjecture, although we do not know if it is correct or not.

#### Conjecture 2.

*In Scenario 4, if any three genes g_i_, g_j_, g_l_ in g*_1_,…, *g_k_ form a common subsequence, then g*_1_,…, *g_k_ form a common subsequence*.

To solve Scenario 4,construct a 3-uniform hypergraph 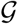 as following [15]: vertices are genes 1,…,*n*; there is a 3-hyperedge (undirected) that links genes *g_i_, g_j_, g_k_* if and only if they form a common subsequence.

#### Proposition 3.

*If Conjecture 2 holds, then finding the longest common sequence in Scenario 4 can be reduced to the maximum clique problem for* 3*-uniform hypergraphs.*

*Proof.* If *g*_1_,…, *g_k_* form a common subsequence, then any three genes *g_i_, g_j_, g_l_* has a 3-hyperedge, and *g*_1_,…, *g_k_* form a complete subgraph. If *g*_1_,…, *g_k_* form a complete subgraph, then any three genes *g_i_, g_j_, g_l_* form a common subsequence. By Conjecture 2, this means *g*_1_,…, *g_k_* form a common subsequence. Therefore, there is a bijection between common subsequence and complete subgraph. If we can find the maximum clique problem for 3-uniform hypergraphs, then it corresponds to the longest common subsequence.

We have reduced Scenario 4 into the maximum clique problem for 3-uniform hypergraphs, which is also NP-hard [77]. There have been some algorithms for the maximum clique problem for 3-uniform hypergraphs [63, 57]. For completeness, we propose a simple idea: Repeatedly delete the gene that has the smallest degree, until we have a complete subgraph that any three genes have a 3-hyperedge that links them. We summarize this greedy heuristic method as Algorithm 6. This algorithm is easy to understand, and can provide some intuition. We do not claim that Algorithm 6 is comparable to other sophisticated algorithms.

We test Algorithm 6 on random graphs. Construct a random graph with *n* genes, and any two genes have probability 0.5 to have an edge in between. Use brute-force search to find the maximum clique, and compare its size with the result of Algorithm 6. For each *n* ≤ 15, we repeat this for 10000 times, and every time Algorithm 6 returns the correct result. Therefore, for small random graphs, the 95% credible interval for the success rate of Algorithm 6 is [0.9997, 1]. We can claim that Algorithm 6 is a good heuristic algorithm that fails with a very small probability. Since finding the true maximum clique requires exponentially slow brute-force search, we do not test on very large graphs.

Nevertheless, Algorithm 6 does not always produce the correct result. See Fig. 7 for a counterexample. Here each gene in 1, 2, 3, 4, 5, 6 has degree 4, while each gene in 7, 8, 9, 10 has degree 3.When applying Algorithm 6, genes 7, 8, 9, 10 are first deleted, and the final result just has three genes, such as (1, 3, 5). However, the longest common subsequence (7, 8, 9, 10) has four genes.

**Figure 7:**
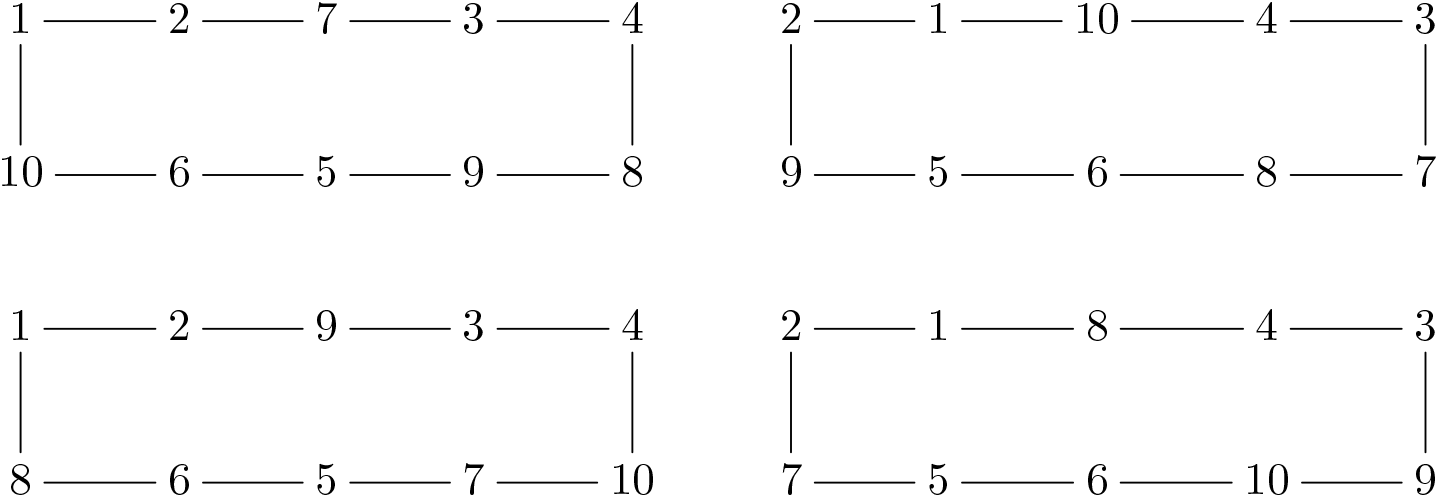
Four circular sequences. The longest common subsequence is (7, 8, 9, 10). This counterexample fails Algorithm 6.

Assume we have *m* sequences with *n* genes. In general, the copy number of a gene is small, and we can assume the length of each sequence is 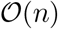. The time complexities of Step 2 and Step 3 in Algorithm 6 are 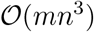 and 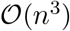, and the overall time complexity is 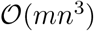. The space complexity is trivially 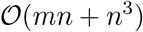.

**Algorithm 6:**
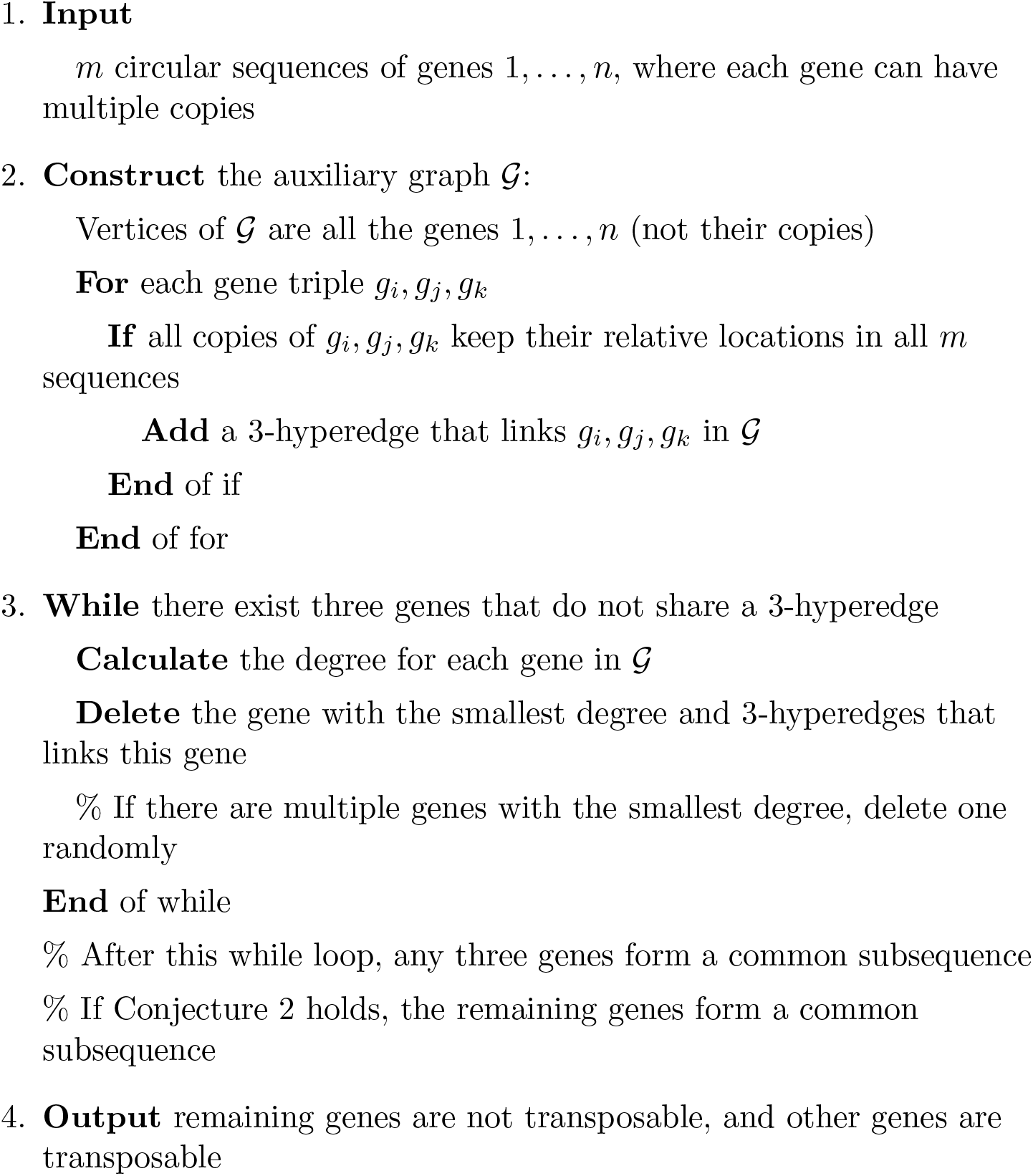
A heuristic method for detecting transposable genes in Scenario 4.

## 7 Discussion

Certain patients of myeloproliferative neoplasm have two mutations, JAK2 V617F and TET2. The temporal order of these two mutations could affect various macroscopic and microscopic features [52]. One explanation for this phenomenon is that mutations can lead to different spatial orders, namely gene transpositions, but different transpositions are not interchangeable [56]. Algorithms in this paper can be used to determine the transposable genes and study how different temporal gene (mutation) orders and spatial gene orders interfere with each other.

A gene *g_i_* might be missing in some sequences. Since *g_i_* is not in any longest common subsequence, it should be a proper-transposable gene. This gene can be directly removed before applying corresponding algorithms.

We can adopt a stricter definition of transposable genes to exclude a gene which only changes its relative position in a few (no more than *l*, where *l* is small enough) sequences. Then we should consider the longest sequence which is a common subsequence of at least *m* – *l* sequences. We can run the corresponding algorithm for every *m* – l sequences. Thus the total time complexity will be multiplied by a factor of *m^l^*.

In Scenario 1 and Scenario 2 (linear/circular sequences without duplicated genes), if each sequence has *n* genes, and the longest common subsequence has length *n* – *k*, then there are at most *k* proper-transposable genes. About quasi-transposable genes, inspired by Lemma 1, we have the following guess.

#### Conjecture 3.

*Consider m linear/circular sequences with n genes without multiple copies. Assume the length of the longest common subsequence is n – k, and there are l proper-transposable genes. Then the number of quasi-transposable genes is no larger than* 2(*k* – *l*).

When *l* + 2(*k* – *l*) ≤ *n*, in both linear and circular scenarios, we can find examples with 2(*k* – *l*) quasi-transposable genes.

## 8 Conclusion

In this paper, we study the problem of determining transposable genes in gene sequences, and design Algorithms 1–6 for different scenarios. To apply those algorithms, one needs to apply genomic annotation tools to transform raw DNA sequencing data into gene sequences, and replace gene names by numbers. Those algorithms have at most *O*(*mn*^3^) time complexity, where *m* is the number of sequences, and *n* is the number of genes. Thus they can run in a reasonable time for most applications. We prove that the latter two scenarios are NP-hard (Propositions 1,2), and propose two unresolved problems (Conjectures 2,3) in discrete mathematics.

We start with gene sequences and determine translocated genes. Therefore, short transposons (possibly shorter than a gene) cannot be determined. Besides, we do not determine specific genomic rearrangement events. We aim at determining which genes are able to translocate. Specifically, we study how many longest common subsequences contain a certain gene, as a measure for its “stability”. This mesoscopic viewpoint can be intriguing for understanding changes in genome.

The results in this paper are not limited to Scenarios 1–4. They can be applied to other bioinformatics situations, or even other fields that need discrete mathematics tools, such as text processing, compiler optimization, data analysis, image analysis [22]. Besides, algorithms in this paper might be able to detect non-syntenic regions [36].

There are some possible future directions: (1) prove Conjectures 2,3; (2) extend Proposition 3 to find more efficient solutions to Scenario 4; (3) determine whether genes appear in all longest common subsequences in other similar scenarios.

## Acknowledgments

The author would like to thank Zhongkai Zhao for helping with designing Algorithm 1. The author would like to thank Lucas Böttcher and anonymous reviewers for providing helpful comments.

## References

[1] Adi, S. S., Braga, M. D., Fernandes, C. G., Ferreira, C. E., Martinez, F. V., Sagot, M.-F., Stefanes, M. A., Tjandraatmadja, C., and Wakabayashi, Y. Repetition-free longest common subsequence. Discret. Appl. Math. 158, 12 (2010), 1315–1324.

[2] Angelini, E., Wang, Y., Zhou, J. X., Qian, H., and Huang, S. A model for the intrinsic limit of cancer therapy: Duality of treatment-induced cell death and treatment-induced stemness. PLOS Comput. Biol. 18, 7 (2022), e1010319.

[3] Babakhani, S., and Oloomi, M. Transposons: the agents of antibiotic resistance in bacteria. J. Basic Microbiol. 58, 11 (2018), 905–917.

[4] Backurs, A., and Indyk, P. Edit distance cannot be computed in strongly subquadratic time (unless SETH is false). In Proceedings of the Forty-Seventh Annual ACM Symposium on Theory of Computing (2015), pp. 51–58.

[5] Bergroth, L., Hakonen, H., and Raita, T. A survey of longest common subsequence algorithms. In Proceedings Seventh International Symposium on String Processing and Information Retrieval. SPIRE 2000 (2000), IEEE, pp. 39–48.

[6] Blanchette,M., Kunisawa, T., and Sankoff, D. Parametric genome rearrangement. Gene 172, 1 (1996), GC11–GC17.

[7] Blum, C., Djukanovic, M., Santini, A., Jiang, H., Li, C.-M., Manyà, F., and Raidl, G. R. Solving longest common subsequence problems via a transformation to the maximum clique problem. Comput. Oper. Res. 125 (2021), 105089.

[8] Bohnenkämper, L., Braga, M. D., Doerr, D., and Stoye, J. Computing the rearrangement distance of natural genomes. J. Comput. Biol. 28, 4 (2021), 410–431.

[9] Brůna, T., Hoff, K. J., Lomsadze, A., Stanke, M., and Borodovsky, M. BRAKER2: automatic eukaryotic genome annotation with GeneMark-EP+ and AUGUSTUS supported by a protein database. NAR Genom. Bioinform. 3, 1 (2021), lqaa108.

[10] Chen, X., and Li, D. ERVcaller: identifying polymorphic endogenous retrovirus and other transposable element insertions using wholegenome sequencing data. Bioinformatics 35, 20 (2019), 3913–3922.

[11] Chen, X., Wang, Y., Feng, T., Yi, M., Zhang, X., and Zhou, D. The overshoot and phenotypic equilibrium in characterizing cancer dynamics of reversible phenotypic plasticity. J. Theor. Biol. 390 (2016), 40–49.

[12] Chen, Z.-Z., Gao, Y., Lin, G., Niewiadomski, R., Wang, Y., and Wu, J. A space-efficient algorithm for sequence alignment with inversions and reversals. Theor. Comput. Sci. 325, 3 (2004), 361–372.

[13] DeNicola, G. M., Karreth, F. A., Adams, D. J., and Wong, C. C. The utility of transposon mutagenesis for cancer studies in the era of genome editing. Genome Biol. 16, 1 (2015), 1–15.

[14] Dessalles, R., Pan, Y., Xia, M., Maestrini, D., D’Orsogna, M. R., and Chou, T. How naive t-cell clone counts are shaped by heterogeneous thymic output and homeostatic proliferation. Front. Immunol. 12 (2021).

[15] Diestel, R. Graph Theory, 5 ed. Springer, Berlin, 2017.

[16] Diop, S. I., Subotic, O., Giraldo-Fonseca, A., Waller, M., Kirbis, A., Neubauer, A., Potente, G., Murray-Watson, R., Boskovic, F., Bont, Z., et al. A pseudomolecule-scale genome assembly of the liverwort marchantia polymorpha. Plant J. 101, 6 (2020), 1378–1396.

[17] Evrony, G. D., Hinch, A. G., and Luo, C. Applications of singlecell dna sequencing. Annu. Rev. Genomics Hum. Genet. 22 (2021), 171.

[18] Gardner, E. J., Lam, V. K., Harris, D. N., Chuang, N. T., Scott, E. C., Pittard, W. S., Mills, R. E., Devine, S. E., Consortium,. G. P., et al. The Mobile Element Locator Tool (MELT): population-scale mobile element discovery and biology. Genome Res. 27, 11 (2017), 1916–1929.

[19] Goel, M., Sun, H., Jiao, W.-B., and Schneeberger, K. Syri: finding genomic rearrangements and local sequence differences from whole-genome assemblies. Genome Biol. 20, 1 (2019), 1–13.

[20] Goubert, C., Craig, R. J., Bilat, A. F., Peona, V., Vogan, A. A., and Protasio, A. V. A beginner’s guide to manual curation of transposable elements. Mob. DNA 13, 1 (2022), 1–19.

[21] Gu, W., Zhang, F., and Lupski, J. R. Mechanisms for human genomic rearrangements. Pathogenetics 1, 1 (2008), 1–17.

[22] Hajiaghayi, M., Seddighin, S., and Sun, X. Massively parallel approximation algorithms for edit distance and longest common subsequence. In Proceedings of the Thirtieth Annual ACM-SIAM Symposium on Discrete Algorithms (2019), SIAM, pp. 1654–1672.

[23] Hirschberg, D. S. A linear space algorithm for computing maximal common subsequences. Commun. ACM 18, 6 (1975), 341–343.

[24] Huang, K., Yang, C.-B., Tseng, K.-T., et al. Fast algorithms for finding the common subsequences of multiple sequences. In Proceedings of the International Computer Symposium (2004), Citeseer, pp. 1006–1011.

[25] Ibal, J. C., Pham, H. Q., Park, C. E., and Shin, J.-H. Information about variations in multiple copies of bacterial 16s rRNA genes may aid in species identification. PLOS ONE 14, 2 (2019), e0212090.

[26] Imbeault, M., Helleboid, P.-Y., and Trono, D. Krab zinc-finger proteins contribute to the evolution of gene regulatory networks. Nature 543, 7646 (2017), 550–554.

[27] Islam, M., Saifullah, C., Asha, Z. T., Ahamed, R., et al. Chemical reaction optimization for solving longest common subsequence problem for multiple string. Soft Comput. 23, 14 (2019), 5485–5509.

[28] Ivics, Z., and Izsvák, Z. The expanding universe of transposon technologies for gene and cell engineering. Mob. DNA 1, 1 (2010), 1–15.

[29] Jiang, D.-Q., Wang, Y., and Zhou, D. Phenotypic equilibrium as probabilistic convergence in multi-phenotype cell population dynamics. PLOS ONE 12, 2 (2017), e0170916.

[30] Jiang, H., Li, C.-M., and Manya, F. Combining efficient preprocessing and incremental MaxSAT reasoning for maxclique in large graphs. In Proceedings of the twenty-second European conference on artificial intelligence (2016), pp. 939–947.

[31] Jiang, T., Lin, G.-H., Ma, B., and Zhang, K. The longest common subsequence problem for arc-annotated sequences. In Annual Symposium on Combinatorial Pattern Matching (2000), Springer, pp. 154–165.

[32] Kang, Y., Gu, C., Yuan, L., Wang, Y., Zhu, Y., Li, X., Luo, Q., Xiao, J., Jiang, D., Qian, M., et al. Flexibility and symmetry of prokaryotic genome rearrangement reveal lineage-associated core-gene-defined genome organizational frameworks. mBio 5, 6 (2014), e01867–14.

[33] Kazazian, H. H., Wong, C., Youssoufian, H., Scott, A. F., Phillips, D. G., and Antonarakis, S. E. Haemophilia A resulting from de novo insertion of L1 sequences represents a novel mechanism for mutation in man. Nature 332, 6160 (1988), 164–166.

[34] Kececioglu, J., and Sankoff, D. Exact and approximation algorithms for sorting by reversals, with application to genome rearrangement. Algorithmica 13, 1 (1995), 180–210.

[35] Kiyomi, M., Horiyama, T., and Otachi, Y. Longest common subsequence in sublinear space. Inf. Process. Lett. 168 (2021), 106084.

[36] Lee, K.-C., and Kim, S.-S. Non-synteny regions in the human genome. Genom. Inform. 8, 2 (2010), 86–89.

[37] Li, C.-M., Jiang, H., and Manyà, F. On minimization of the number of branches in branch-and-bound algorithms for the maximum clique problem. Comput. Oper. Res. 84 (2017), 1–15.

[38] Lin, C.-H., Lian, C.-Y., Hsiung, C. A., and Chen, F.-C. Changes in transcriptional orientation are associated with increases in evolutionary rates of enterobacterial genes. BMC Bioinform. 12, 9 (2011), 1–8.

[39] Maier, D. The complexity of some problems on subsequences and supersequences. J. ACM 25, 2 (1978), 322–336.

[40] Makałowski, W., Gotea, V., Pande, A., and Makałowska, I. Transposable elements: Classification, identification, and their use as a tool for comparative genomics. In Evolutionary Genomics. Springer, 2019, pp. 177–207.

[41] McClintock, B. The origin and behavior of mutable loci in maize. Proc. Natl. Acad. Sci. U.S.A. 36, 6 (1950), 344–355.

[42] Mills, R. E., Bennett, E. A., Iskow, R. C., and Devine, S. E. Which transposable elements are active in the human genome? Trends Genet. 23, 4 (2007), 183–191.

[43] Mitsuhashi, S., Ohori, S., Katoh, K., Frith, M. C., and Matsumoto, N. A pipeline for complete characterization of complex germline rearrangements from long dna reads. Genome Med. 12, 1 (2020), 1–17.

[44] Mousavi, S. R., and Tabataba, F. An improved algorithm for the longest common subsequence problem. Comput. Oper. Res. 39, 3 (2012), 512–520.

[45] Mustajoki, S., Ahola, H., Mustajoki, P., and Kauppinen, R. Insertion of Alu element responsible for acute intermittent porphyria. Hum. Mutat. 13, 6 (1999), 431–438.

[46] Ngomade, A. N., Myoupo, J. F., and Tchendji, V. K. A dominant point-based parallel algorithm that finds all longest common subsequences for a constrained-mlcs problem. J. Comput. Sci. 40 (2020), 101070.

[47] Niu, X.-M., Xu, Y.-C., Li, Z.-W., Bian, Y.-T., Hou, X.-H., Chen, J.-F., Zou, Y.-P., Jiang, J., Wu, Q., Ge, S., et al. Transposable elements drive rapid phenotypic variation in Capsella rubella. Proc. Natl. Acad. Sci. U.S.A. 116, 14 (2019), 6908–6913.

[48] Niu, Y., Wang, Y., and Zhou, D. The phenotypic equilibrium of cancer cells: From average-level stability to path-wise convergence. J. Theor. Biol. 386 (2015), 7–17.

[49] Noorani, I., Bradley, A., and de la Rosa, J. CRISPR and transposon in vivo screens for cancer drivers and therapeutic targets. Genome Biol. 21, 1 (2020), 1–22.

[50] Nowacki, M., Higgins, B. P., Maquilan, G. M., Swart, E. C., Doak, T. G., and Landweber, L. F. A functional role for transposases in a large eukaryotic genome. Science 324, 5929 (2009), 935–938.

[51] Orozco-Arias, S., Piña, J. S., Tabares-Soto, R., Castillo-Ossa, L. F., Guyot, R., and Isaza, G. Measuring performance metrics of machine learning algorithms for detecting and classifying transposable elements. Processes 8, 6 (2020), 638.

[52] Ortmann, C. A., Kent, D. G., Nangalia, J., Silber, Y., Wedge, D. C., Grinfeld, J., Baxter, E. J., Massie, C. E., Papaemmanuil, E., Menon, S., et al. Effect of mutation order on myeloproliferative neoplasms. N. Engl. J. Med. 372, 7 (2015), 601–612.

[53] Payer, L. M., and Burns, K. H. Transposable elements in human genetic disease. Nat. Rev. Genet. 20, 12 (2019), 760–772.

[54] Rahrmann, E. P., Collier, L. S., Knutson, T. P., Doyal, M. E., Kuslak, S. L., Green, L. E., Malinowski, R. L., Roethe, L., Akagi, K., Waknitz, M., et al. Identification of PDE4D as a proliferation promoting factor in prostate cancer using a Sleeping Beauty transposon-based somatic mutagenesis screen. Cancer Res. 69, 10 (2009), 4388–4397.

[55] Reneker, J., Lyons, E., Conant, G. C., Pires, J. C., Freeling, M., Shyu, C.-R., and Korkin, D. Long identical multispecies elements in plant and animal genomes. Proc. Natl. Acad. Sci. U.S.A. 109, 19 (2012), E1183–E1191.

[56] Roquet, N., Soleimany, A. P., Ferris, A. C., Aaronson, S., and Lu, T. K. Synthetic recombinase-based state machines in living cells. Science 353, 6297 (2016), aad8559.

[57] Rota Bulò, S., and Pelillo, M. A continuous characterization of maximal cliques in k-uniform hypergraphs. In International conference on learning and intelligent optimization (2007), Springer, pp. 220–233.

[58] Rowley, M. J., and Corces, V. G. Organizational principles of 3D genome architecture. Nat. Rev. Genet. 19, 12 (2018), 789–800.

[59] Salzberg, S. L. Next-generation genome annotation: we still struggle to get it right, 2019.

[60] Sha, Y., Wang, S., Zhou, P., and Nie, Q. Inference and multiscale model of epithelial-to-mesenchymal transition via single-cell transcriptomic data. Nucleic Acids Res. 48, 17 (2020), 9505–9520.

[61] Terauds, V., and Sumner, J. Maximum likelihood estimates of rearrangement distance: implementing a representation-theoretic approach. Bull. Math. Biol. 81, 2 (2019), 535–567.

[62] Tesler, G. Efficient algorithms for multichromosomal genome rearrangements. J. Comput. Syst. Sci. 65, 3 (2002), 587–609.

[63] Torres-Jimenez, J., Perez-Torres, J. C., and Maldonado-Martinez, G. hClique: An exact algorithm for maximum clique problem in uniform hypergraphs. Discrete Math. Algorithms Appl. 9, 06 (2017), 1750078.

[64] Valiente, G. Algorithms on Trees and Graphs. Springer, Berlin, 2002.

[65] Verma, S. C., Qian, Z., and Adhya, S. L. Architecture of the Escherichia coli nucleoid. PLOS Genet. 15, 12 (2019), e1008456.

[66] Wang, Q., Korkin, D., and Shang, Y. A fast multiple longest common subsequence (MLCS) algorithm. IEEE Trans. Knowl. Data Eng. 23, 3 (2010), 321–334.

[67] Wang, T., Weiss, A., Aqeel, A., Wu, F., Lopatkin, A. J., David, L. A., and You, L. Horizontal gene transfer enables programmable gene stability in synthetic microbiota. Nat. Chem. Biol. (2022), 1–8.

[68] Wang, X., Wang, L., and Zhu, D. Efficient computation of longest common subsequences with multiple substring inclusive constraints. J. Comput. Biol. 26, 9 (2019), 938–947.

[69] Wang, Y. Some Problems in Stochastic Dynamics and Statistical Analysis of Single-Cell Biology of Cancer. Ph.d. thesis, University of Washington, 2018.

[70] Wang, Y. Two metrics on rooted unordered trees with labels. Algorithms Mol. Biol. 17, 1 (2022), 1–17.

[71] Wang, Y., Cai, S., and Yin, M. Two efficient local search algorithms for maximum weight clique problem. In Thirtieth AAAI Conference on Artificial Intelligence (2016), pp. 805–811.

[72] Wang, Y., Kropp, J., and Morozova, N. Biological notion of positional information/value in morphogenesis theory. Int. J. Dev. Biol. 64, 10-11-12 (2020), 453–463.

[73] Wang, Y., and Wang, L.Causal inference in degenerate systems: An impossibility result. In International Conference on Artificial Intelligence and Statistics (2020), PMLR, pp. 3383–3392.

[74] Wang, Y., and Wang, Z. Inference on the structure of gene regulatory networks. J. Theor. Biol. 539 (2022), 111055.

[75] Wang, Y., Zhang, B., Kropp, J., and Morozova, N. Inference on tissue transplantation experiments. J. Theor. Biol. 520 (2021), 110645.

[76] Wei, S., Wang, Y., Yang, Y., and Liu, S. A path recorder algorithm for Multiple Longest Common Subsequences (MLCS) problems. Bioinformatics 36, 10 (2020), 3035–3042.

[77] Wu, Q., and Hao, J.-K. A review on algorithms for maximum clique problems. Eur. J. Oper. Res. 242, 3 (2015), 693–709.

[78] Xia, M., Greenman, C. D., and Chou, T. PDE models of adder mechanisms in cellular proliferation. SIAM J. Appl. Math. 80, 3 (2020), 1307–1335.

[79] Yu, T., Huang, X., Dou, S., Tang, X., Luo, S., Theurkauf, W. E., Lu, J., and Weng, Z. A benchmark and an algorithm for detecting germline transposon insertions and measuring de novo transposon insertion frequencies. Nucleic Acids Res. 49, 8 (2021), e44–e44.

[80] Zhou, D., Wang, Y., and Wu, B. A multi-phenotypic cancer model with cell plasticity. J. Theor. Biol. 357 (2014), 35–45.

[81] Zhou, P., Wang, S., Li, T., and Nie, Q. Dissecting transition cells from single-cell transcriptome data through multiscale stochastic dynamics. Nat. Commun. 12, 1 (2021), 1–15.

[82] Zhou, W., Liang, G., Molloy, P. L., and Jones, P. A. Dna methylation enables transposable element-driven genome expansion. Proc. Natl. Acad. Sci. U.S.A. 117, 32 (2020), 19359–19366.

[83] Zimin, A. V., Puiu, D., Luo, M.-C., Zhu, T., Koren, S., Marçais, G., Yorke, J. A., Dvořák, J., and Salzberg, S. L. Hybrid assembly of the large and highly repetitive genome of aegilops tauschii, a progenitor of bread wheat, with the masurca mega-reads algorithm. Genome Res. 27, 5 (2017), 787–792.

